# Genetic rescue without genomic swamping in wild populations

**DOI:** 10.1101/701706

**Authors:** Sarah W. Fitzpatrick, Gideon S. Bradburd, Colin T. Kremer, Patricia E. Salerno, Lisa M. Angeloni, W. Chris Funk

## Abstract

Gene flow is an enigmatic evolutionary force because it can limit adaptation but can also help populations escape inbreeding depression. Manipulating gene flow for conservation purposes is a controversial, but potentially powerful management strategy. We use multigenerational pedigrees and genomics to test demographic and evolutionary consequences of manipulating gene flow in two isolated wild Trinidadian guppy populations. We found that on average, hybrids lived longer and reproduced more. Despite overall genome-wide homogenization, alleles potentially associated with local adaptation were not entirely swamped by gene flow. Our results suggest that combining new genomic variation from immigrants with potentially adaptive variation from the recipient population resulted in highly fit hybrids and subsequent increases in population size. Contrary to the prevailing view that gene flow constrains adaptation, our study shows that immigration can produce long-term fitness benefits in small populations without swamping locally adaptive variation.

## Introduction

Gene flow is a fundamental force in evolutionary biology; it determines the distribution of genetic variation exposed to natural selection, can inhibit or facilitate local adaptation, and contributes to species cohesion (*1, 2*). Gene flow is also of fundamental concern in conservation biology, as it can affect the capacity of a population to escape inbreeding depression or to adapt to rapidly changing environmental conditions (*3*). Altered gene flow patterns, due to natural or anthropogenic processes, can have major demographic consequences. Too much gene flow can cause population declines or extirpation via disruption of local adaptation (*4*), outbreeding depression (*5*), or invasive hybridization (*6*). However, too little gene flow can result in fixation of deleterious alleles in small and isolated populations, potentially also leading to their decline or extirpation (*7*). Predicting evolutionary and demographic effects of altered gene flow patterns is thus a fundamental goal in evolutionary ecology and a major challenge for biodiversity conservation.

Debates over the net effect and context dependence of gene flow on population fitness has real-world consequences for the conservation and management of vulnerable populations. Assisted gene flow, the intentional translocation of individuals, is a potentially powerful tool for counteracting inbreeding depression and/or mitigating maladaptation due to climate change (*8*). ‘Genetic rescue’, a process by which the introduction of new alleles causes an increase in population growth (*9, 10*), has been tested experimentally in controlled lab and greenhouse environments (*11*-*13*), and observed in the wild in a handful of iconic examples of human-assisted gene flow for conservation purposes (*14, 15*). Despite increasing evidence that genetic rescue is often a plausible outcome, concerns over outbreeding depression and biotic homogenization limit its use to cases of last resort.

Although there is general consensus that outcrossing inbred populations provides initial fitness benefits (*16*), limited understanding of the mechanisms and long-term effects of gene flow on fitness constrains the use of assisted gene flow as a tool for management of imperiled populations. Most studies of genetic rescue in the wild are correlative, unreplicated, and span one or two generations (*10*). In later generations, initial fitness benefits from immigrant × resident crosses may break down when deleterious interactions between homozygous loci become exposed (i.e., Dobzhansky-Muller incompatibilities). Another concern is the genomic swamping of locally adaptive variation (*6, 17*). Small populations may be especially vulnerable to genetic swamping from large immigrant populations due to differences in effective population size and genetic load (*18*). Thus, our understanding of the demographic and evolutionary impacts of gene flow into isolated populations in nature is still lacking experimental elements such as: (i) replicated, multigenerational tests of gene flow manipulations in wild populations, which may yield different conclusions than lab experiments, and (ii) genomic data from genetic rescue studies to test the extent of genomic swamping or maintenance of locally adaptive variation.

We took advantage experimental translocations of Trinidadian guppy populations to test the demographic and evolutionary consequences of new gene flow into small, isolated populations in the wild. Trinidadian guppies (*Poecilia reticulata*) are a model system for studying contemporary evolution, ecology, and conservation (*19*). Here, replicated translocations of guppies from a downstream site into headwater sites upstream from native, recipient guppy populations ((*20*); Fig. 1A) provided the opportunity to study the demographic and evolutionary effects of gene flow. Recipient and source populations came from environments with different predation and resource regimes and thus were presumed to be adaptively differentiated (*21*). Previous work on these same populations suggests genetic rescue occurred as a result of the translocations: gene flow into recipient populations was accompanied by large increases in population size, which were attributed to high recruitment rates in immigrant and hybrid ancestry classes (*22*), while locally favored traits were retained (*23*). In this study we used multigenerational wild pedigrees and genomic data to determine the mechanisms—from genes to individual fitness—that caused genetic rescue. We addressed three specific questions:

*(i)* Is genetic rescue caused primarily by superior immigrants that have higher fitness than residents, or by highly fit hybrids?
*(ii)* What are the genomic consequences of new gene flow?
*(iii)* Is locally adaptive genetic variation maintained by selection in the face of gene flow?

**Fig. 1.**
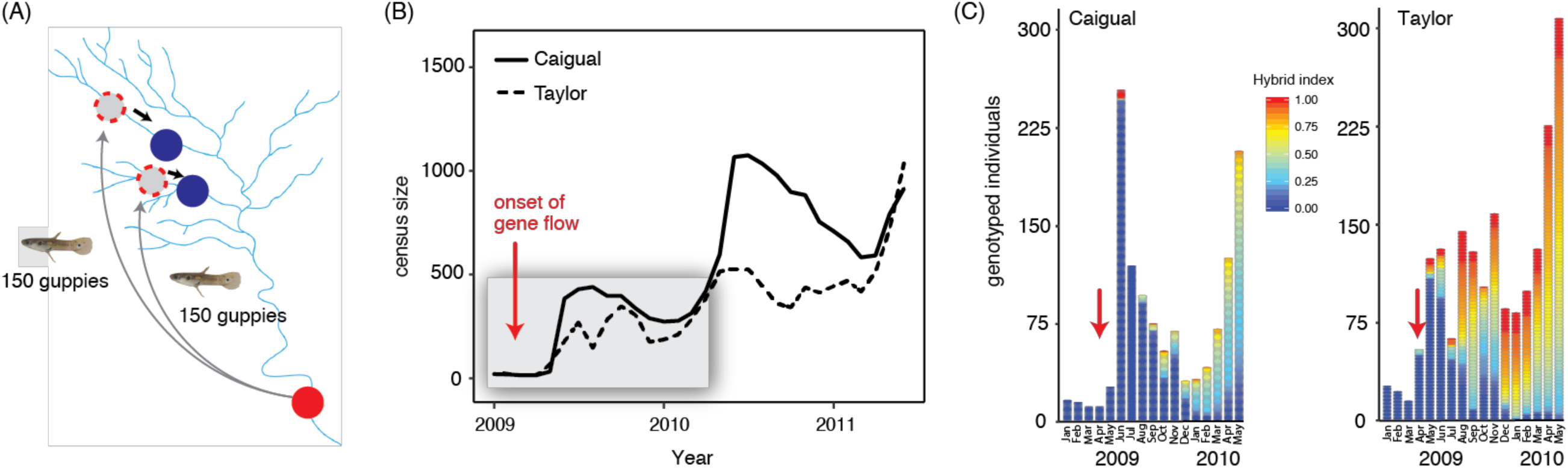
Gene flow manipulation experiments in Trinidad. (A) Map of the Guanapo River drainage. In 2009, guppies were translocated from a downstream high predation locality (red) into two headwater sites (dashed red) that were upstream of native recipient populations in low predation environments (dark blue). Unidirectional, downstream gene flow began shortly after the introductions, indicated by black arrows. (B) Census sizes in Caigual (solid) and Taylor (dashed) following the onset of gene flow from the upstream introduction sites. Grey box indicates the timespan in which all captured individuals were genotyped at 12 microsatellite loci. (C) Temporal patterns of continuous hybrid index assignments throughout the first seventeen months of the study (∼four to six guppy generations). Recipient populations prior to gene flow had a hybrid index = 0 and pure immigrant individuals had a hybrid index = 1. Hybrid indices were assigned using data from 12 microsatellite loci. Red arrows indicate the onset of gene flow.

To answer these questions, we combined long-term individual-based capture-mark-recapture and genetic monitoring with pedigree reconstruction. We evaluated the effect of individual hybrid index on major fitness components (longevity and total lifetime reproductive success). We also used genome-wide restriction site-associated DNA sequencing (RADseq) to examine the extent to which the genome became homogenized by gene flow and to test whether alleles associated with the local environment were maintained at a higher frequency than neutral loci in the face of high gene flow. The integration of these different data types allowed us to dissect in unprecedented detail the ecological and evolutionary mechanisms of genetic rescue.

## Results

We captured, uniquely marked, and monitored 9,590 Trinidadian guppies in focal recipient populations in the Caigual and Taylor Rivers, located on the south slope of the Northern Range Mountains in Trinidad, over the course of 29 consecutive months (∼eight to ten guppy generations). Detection probabilities were high in both streams with monthly averages of 0.83 in Taylor and 0.86 in Caigual (*22*). The first three capture events occurred prior to upstream translocations. Microsatellite genotypes for every individual captured during the first 17 months of the study (∼four to six guppy generations) were used to reconstruct pedigrees (n=2831) and assign hybrid indices ranging from 0 (pure recipient genotype) to 1 (pure immigrant genotype). We captured 63 fish with immigrant genotypes in the Caigual recipient population and 753 immigrant genotypes in the Taylor recipient population.

### Sustained population growth

Following the onset of gene flow, population sizes increased nearly ten-fold throughout the two years in which Caigual and Taylor recipient populations were censused ((*22*); Fig. 1B), with modest fluctuations driven by typical wet/dry season dynamics (*24*). Prior to upstream translocations, Caigual and Taylor populations were composed entirely of pure recipient genotypes (Fig. 1C). Immigration led to an increase in the frequency of hybrid genotypes throughout the study’s duration. By the end of the 17-month period for which we had individual genotype data, both populations were composed mostly of hybrid and immigrant individuals.

### High hybrid fitness

Hybrid index was a strong predictor of variation in fitness in both streams. Hybrids and/or pure immigrants lived longer (Fig. 2A-B) and had higher lifetime reproductive success (Fig. 2C-D) than pure recipient individuals. Quadratic models relating fitness to hybrid index consistently outperformed linear or constant models. In some cases, these quadratic relationships clearly showed that hybrid genotypes lived longer (e.g., males in Caigual and both sexes in Taylor, Fig. 2A-B) and had higher reproductive success (hybrids in Taylor, Fig. 2D) than individuals with pure recipient or immigrant genotypes. In other cases, hybrids and pure immigrant individuals may have had comparable success (Fig. 2C; females in Fig. 2A). We found some evidence of zero-inflation in longevity (i.e., more fish failed to survive beyond their initial capture than expected given the negative binomial distribution) in both Taylor and Caigual. In Caigual, zero-inflation did not appear to vary between sexes or with hybrid index (Table S2, S5). However, in Taylor, zero-inflation peaked at intermediate hybrid indices and was more common among females (Fig. S4). We also found significant zero-inflation in lifetime reproductive success of Taylor fish (i.e., more fish failed to reproduce than expected given the negative binomial distribution), especially those with recipient genotypes (indicated by significant zero-inflation peaking at low hybrid indices; Fig. S4).

**Fig. 2.**
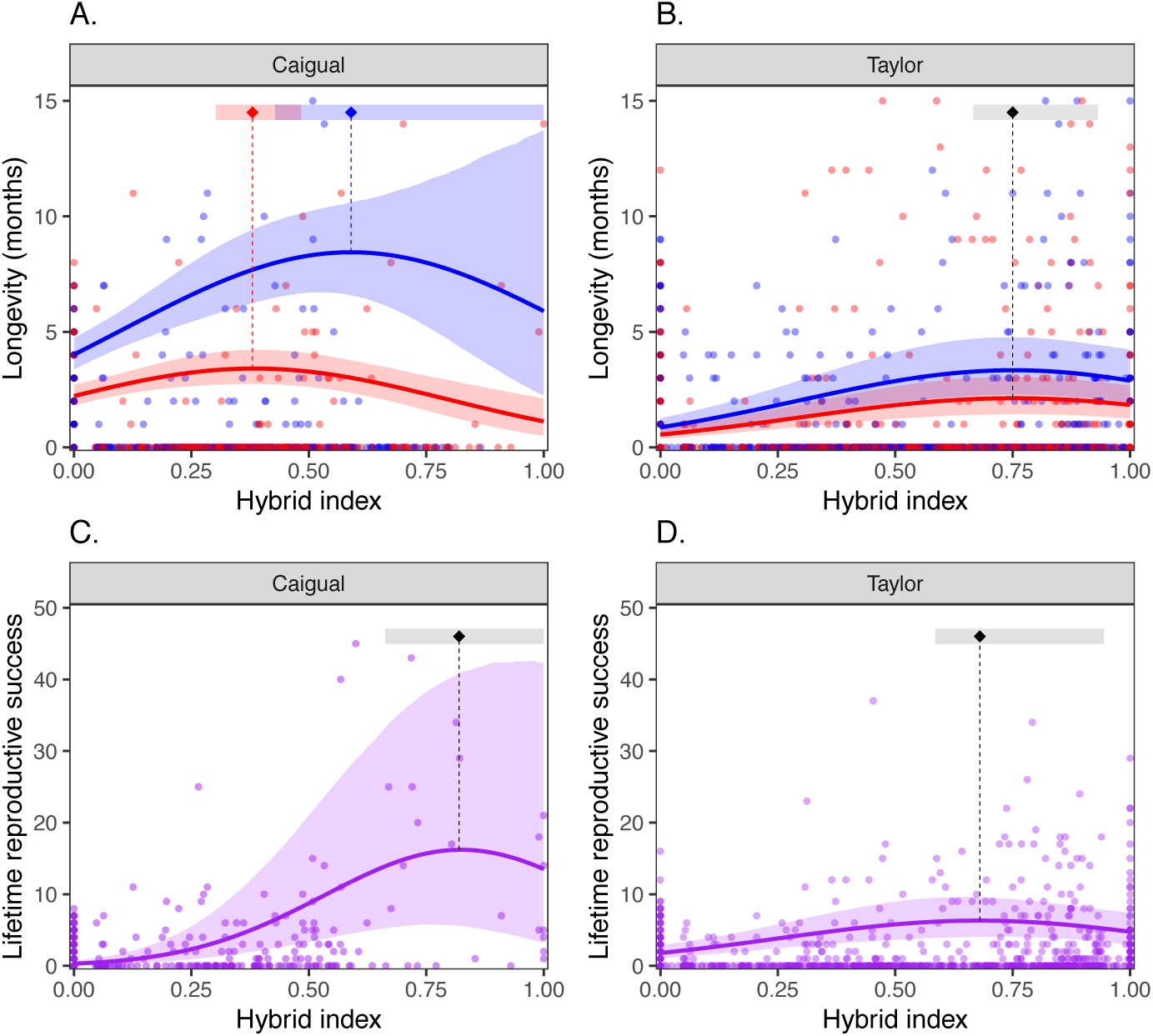
Relationships between hybrid index and fitness. Fitness metrics (longevity and total lifetime reproductive success, LRS) varied quadratically with hybrid index (0 = pure recipient genotype, 1 = pure immigrant genotype). Maxima of the quadratic functions are indicated by vertical dashed lines/diamonds; uncertainty in their positions is indicated by (horizontal) 95% confidence bars. Shading around regression lines displays approximate 95% confidence bands obtained through simulation. Model details appear in Supplemental Tables S1-S5. Longevity differed between males (red) and females (blue) (**A** and **B**). Generally, females lived longer than males, and fish in Caigual lived longer than those in Taylor. In Taylor, male and female longevity had quadratic relationships with hybrid index that differed in magnitude, but peaked at the same location; this differed by sex in Caigual (**A** vs. **B**). LRS varied quadratically with hybrid index and this trend did not differ between males and females (**C** and **D**). Individuals from Taylor generally had lower LRS, and were more likely to not reproduce at all (i.e., we detected significant zero inflation, Fig. S4), especially those with recipient genotypes (hybrid indices near zero). Plotted regressions in panels (**A-B**) and (**D**) display the average predicted success of individuals after controlling for zero inflation. LRS in Caigual individuals (**C**) showed no strong evidence of zero inflation.

### Increased diversity and homogenized variation at most loci

RADseq genotyping of 12,407 SNPs (an average of one locus per 58,542 bp) in pre- and post-gene flow Caigual and Taylor populations and the source population revealed increased genomic variation within recipient populations and substantial genomic homogenization among populations. Before gene flow, the recipient populations were highly distinct from each other and from the source population (dark blue vs. red in Fig. 3A) and showed extremely low levels of genomic variation (Fig. 3B). Genomic differentiation between recipient and source populations decreased dramatically after the onset of gene flow (light blue vs. red in Fig. 3A). Genome-wide average *F*_st_ between the recipient and source populations decreased from 0.29 to 0.01 in Caigual and from 0.31 to 0.02 in Taylor. Recipient Caigual and Taylor populations showed nearly entirely homozygous genomes and extremely low nucleotide diversity, followed by substantial increases in both metrics after the onset of gene flow (Fig. 3B). Ninety-five percent of SNPs were monomorphic in Caigual and 96 % in Taylor prior to gene flow, compared to 22 % and 24 % monomorphic loci after gene flow. Average nucleotide diversity (*π*) increased from 0.01 to 0.22 in Caigual and from 0.01 to 0.21 in Taylor.

**Fig. 3.**
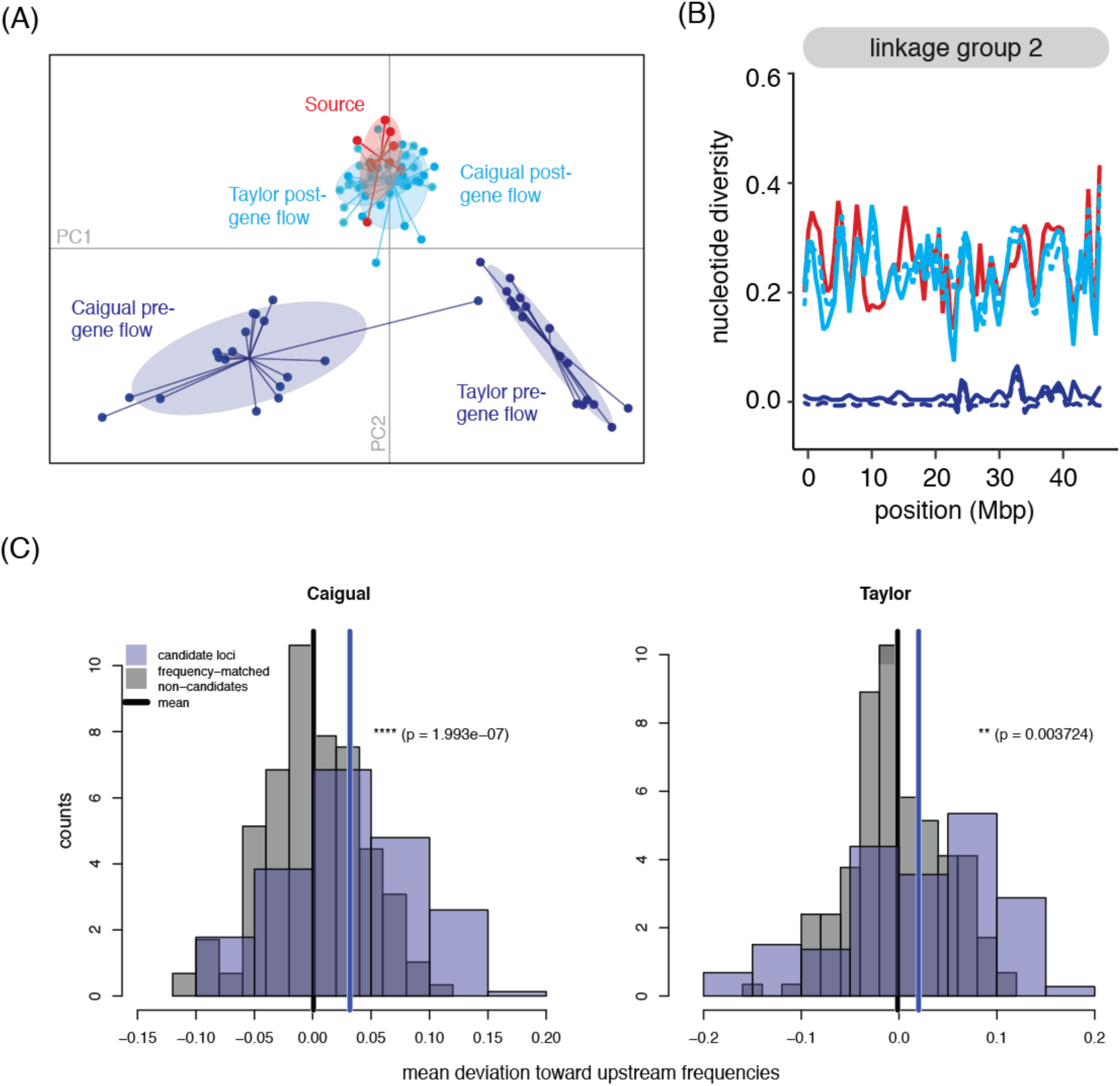
**Genomic consequences of gene flow**. New gene flow caused overall genomic homogenization but candidate adaptive alleles were maintained at higher than expected frequencies. (**A**) PCA plot showing overall population differentiation based on polymorphic SNP loci from the RADseq data. (**B**) Comparison of nucleotide diversity patterns along linkage group two among pre-gene flow (dark blue) and post-gene flow (light blue) Caigual (solid) and Taylor (dashed) populations and the introduction source (red). Similar patterns were found across all 23 linkage groups. (**C**) Distributions of ancestry-polarized deviations in candidate loci vs. frequency-matched non-candidates for both populations. In each stream, the allele frequencies of the candidate loci were significantly closer to the headwater ancestral frequency compared to a set of frequency-matched non-candidates.

### Locally adaptive variation maintained

To study the maintenance of locally adaptive variation, we first needed to identify alleles involved in adaptation to headwater, low predation (LP) environments. We assumed such alleles were favored in both LP recipient populations, yet selected against in the downstream, high predation (HP) population. Using an arbitrary frequency difference cutoff of 0.9 between upstream and downstream populations, we identified a set of 146 such loci spanning all 23 linkage groups that were strong candidates for alleles involved in local adaptation (or linked to putatively adaptive alleles) to the LP environment. An excess of pre-gene flow ancestry at these loci (vs. neutral expectations) in the post-gene flow populations would be evidence of selection for the maintenance of locally adaptive variation in the face of gene flow.

Despite overall genomic homogenization, we found evidence for selective maintenance of alleles at these candidate loci in Caigual and Taylor post-gene flow populations. Using simulations, we inferred that the frequencies of this set of candidate alleles were significantly more similar to the inferred ancestral LP population allele frequencies than expected compared to frequency-matched non-candidate loci (Fig. 3C). In other words, an excess of pre-gene flow ancestry at candidate adaptive loci in post-gene flow populations is consistent with selection for the maintenance of locally adaptive variation that was present in pre-gene flow recipient populations. BLAST query results against the Trinidadian guppy genome showed that 129 out of the 146 candidate loci were located in a gene or within 10 kb of a gene. There was no intersection between these BLAST hits and a set of 40 genes previously identified (*25*) as potential contributors to divergent guppy phenotypes such as growth, vision, and pigment pattern development.

## Discussion

Gene flow between adaptively differentiated populations is typically assumed to swamp local adaptation and reduce fitness, but few studies have mechanistically tested multigenerational fitness effects of gene flow into small and isolated populations in the wild. We documented high hybrid fitness resulting in sustained population growth over multiple generations in replicated populations of Trinidadian guppies. Contrary to the prediction that small populations are especially vulnerable to genomic swamping, we showed that some portions of the recipient genome (associated with the local environment) were maintained, suggesting that genetic load was reduced without compromising potentially important adaptive variation.

Empirical tests of the phenotypic and fitness effects of gene flow in wild populations tend to yield idiosyncratic responses, giving rise to the prevailing wisdom that phenotypic effects of gene flow are trait-specific and net fitness effects are difficult to predict (*3*). An advantage of our design over that of previous genetic rescue studies is replication across two neighboring headwater populations. We observed similar fitness and genomic responses to gene flow in both populations, suggesting that the demographic and evolutionary responses to gene flow under similar conditions are not inherently idiosyncratic. This has important conservation implications: once the relevant factors (i.e., within and among population genetic and phenotypic characteristics) are understood and assessed, the design of successful assisted gene flow programs for threatened populations may be more feasible than expected.

Our pedigree analysis included fish up to six generations following gene flow and we found that fish with intermediate hybrid indices had the highest longevity and lifetime reproductive success (LRS). Given that guppies breed year-round and have overlapping generations, it is difficult to estimate generation-by-generation fitness metrics. However, individuals with a hybrid index between 0.6-0.8 had the highest fitness in all cases except for male longevity in Caigual (fitness peak at 0.35), suggesting that high hybrid fitness extended beyond heterosis in the F_1_ generation (hybrid index = 0.5). Interestingly, the average hybrid index of individuals with maximal fitness (0.8 in Caigual; 0.7 in Taylor) was similar to the average genome-wide hybrid index sampled at the end of our study, 8−10 generations after gene flow, suggesting individuals with hybrid genomes continued to contribute disproportionately to the observed increases in population size.

The extent to which adaptation in small populations is limited by genetic drift and facilitated by occasional gene flow is a question in need of more attention. In our study, recipient populations underwent substantial genomic changes following gene flow, generally conforming to expectations that gene flow increases genetic variation within populations and homogenizes differentiation among populations. However, whereas microsatellite analysis revealed genetic swamping by the immigrant genotype (*22*), genome-wide analysis revealed maintenance of presumably adaptive alleles at higher than expected frequencies. This result suggests that gene flow into small populations does not inevitably swamp locally important variation, and highlights a major advance in the use of genomic data to untangle the complexities of hybridized wild populations (*26*). We note that although candidate adaptive allele frequencies resisted introgression more than our neutral expectations, they did undergo substantial shifts towards the source population. Unlike the single pulse of immigration typically implemented when assisted gene flow is used for rescuing small populations, rates of gene flow in this study were high (93 migrants per generation on average) and continuous. Under the lower rates of migration recommended for assisted gene flow in management, strong selection would likely maintain adaptive alleles at higher frequency than we observed.

We do not yet know the functional significance of alleles that resisted introgression. The lack of BLAST hits to genes implicated in divergently adapted traits was unsurprising given the sparseness of our genotyping across the genome (*27*) and limited understanding of the genomic architecture of local adaptation in guppies. Our candidate loci might be located in (or linked to) relevant genes whose functions are unknown, or they could affect uncharacterized traits involved in local adaption to the headwater environment (e.g., physiological and metabolic traits) that have not been mapped. Given these limitations, we emphasize that it is the signature of selection in the face of such high migration that is itself interesting. Further investigation of differential rates of introgression throughout the genome with higher resolution genomic data will help identify the genomic architecture of local adaptation to headwater environments in guppies. This task will be additionally strengthened by directly linking variable patterns of introgression to changes in traits and individual fitness.

In our view, the scenario studied here represents an ideal management outcome in which gene flow into small, inbred populations causes substantial increases in genomic variation, individual fitness, and population size, but does not wipe out variation presumed to be locally adaptive. To what extent this scenario translates to other organisms, including species of conservation concern, is unknown. However, our results agree with a growing body of literature supporting the idea that gene flow from a closely related source into small, genetically depauperate populations can produce substantial demographic benefits (*12, 13, 28, 29*). These studies support a proposed paradigm shift in the genetic management of small populations from the current default of inaction to a new policy that considers restoring gene flow to recently fragmented populations (*30*). Selection alone may be unlikely to counteract maladaptation caused by rapid global change, especially in small populations with low genetic variation (*31*). In these cases, gene flow may be essential for providing the necessary variation for populations to persist and adapt to fast-paced environmental change.

## Materials and Methods

### Capture-mark-recapture

Detailed mark-recapture methods are provided in (*22*). Briefly, focal stream reaches in the Caigual and Taylor Rivers located on the south slope of the Northern Range Mountains in Trinidad were sampled every month from January 2009 to June 2011. Three sample occasions occurred prior to upstream introductions (see (*20*) for detailed description of the upstream guppy translocation experiment), followed by 26 additional sample occasions. All guppies greater than 14 mm were caught using nets or minnow traps and transported to the lab in Nalgene^®^ (Rochester, NY, USA) bottles filled with stream water. In the lab, fish were housed in aerated tanks separated by location and sex. Fish were anesthetized with dilute MS-222 and processed under a dissecting microscope. New recruits had three scales removed and dried for DNA extraction and were given a unique set of visible implant elastomer marks (Northwest Marine Technologies, Inc., Shaw Island, WA, USA) using eight marking sites and 12 possible colors. The set of marks used in our study was distinct from the set used in the upstream translocations. All fish each month were weighed, photographed, and returned to their exact location within the focal stream reach one to two days after initial capture. Lab mortality was less than 0.5%. In total, we captured and uniquely marked 9,590 individual guppies throughout 29 capture occasions (4,880 in Caigual; 4,710 in Taylor).

### Microsatellite genotyping and pedigree reconstruction

We extracted genomic DNA from scale samples from all individuals caught in the first 17 (of 29) capture occasions using Gentra Puregene Tissue Kits (Qiagen). Guppies were genotyped at 12 microsatellite markers as described in (*22*). Briefly, we amplified loci using Qiagen Type-It Microsatellite Multiplex PCR kits. PCR products combined with HiDi formamide and LIZ size standard were sent to the Life Sciences Core Laboratory at Cornell University to be read on an ABI 3730xl automated sequencer. Genotypes were visualized and scored using the microsatellite plug-in with GENEIOUS 7.1.7 (*32*). We scored two positive controls and one negative control on each plate and found low genotyping error rate (<0.5%). In total, we genotyped 3,298 guppies (1,491 from Caigual; 1,807 from Taylor) at 12 microsatellite loci. We calculated a continuous hybrid index between 0 (“pure” recipient genotype) and 1 (“pure” immigrant genotype) for all individuals using the maximum likelihood method of (*33*) in the GenoDive software (*34*). A set of 20 pre-gene flow fish from Caigual and Taylor and 20 fish from the downstream source were used as reference populations to estimate hybrid index.

We reconstructed wild pedigrees of the Caigual and Taylor focal populations using Colony2 (*35*), specifying a polygamous mating system without inbreeding, a genotyping error rate of 0.005, and the full-likelihood analysis method with “high” likelihood precision and “medium” runs that were repeated five times with different random seeds to maximize correct parentage assignment. Parent-offspring relationships were only assigned if the same assignment was made in at least four out of the five runs. Separate analyses were carried out for each stream and for each successive cohort of new recruits. The set of possible parents for a given sample occasion were all guppies caught during previous occasions and the set of possible offspring were all new recruits to the population. The final pedigree consisted of 1,106 individuals in Caigual (458 maternal links, 655 paternal links) and 1,725 individuals in Taylor (975 maternal links, 994 paternal links) spanning 4-6 overlapping generations.

### Fitness estimates and GLMMs

We estimated two components of total fitness: longevity and total lifetime reproductive success. Longevity was defined as the maximum number of months an individual was confirmed as present in our study (after the first occurrence) using the full mark-recapture dataset. This value is likely an underestimate of true survival due to imperfect detection probability and because new recruits had to be greater than 14 mm (approximately two months old) to be detected. Lifetime reproductive success was defined as the total number of offspring assigned to an individual using the reconstructed pedigree from individuals caught and genotyped in the first 17 months of data collection. We restricted our estimates to individuals captured in the first 13 months because individuals sampled after this cohort lacked enough time to reproduce and have their offspring identified (Fig. S7-8).

We used generalized linear mixed models (GLMMs) to examine the relationship between an individual’s fitness components and their hybrid index, accounting for the month the individual was first captured (i.e., its cohort) as a random effect, and checking for potential effects of sex and zero-inflation. Variation in both fitness components was modelled using a negative binomial distribution; this discrete distribution is appropriate for potentially over-dispersed count data (such as the number of offspring produced) as well as our discretized estimate of lifespan (here, number of months). We considered a suite of competing models describing changes in the negative binomial distribution’s mean as a function of hybrid index and/or sex, using a log link function (Table S1, S4). A subset of these models also tested for zero-inflation (i.e., zeros in excess of those inherently predicted by the negative binomial distribution) and examined whether the extent of zero-inflation varied with hybrid index and/or sex, using a logit link function. All models were fit using the glmmTMB package (*36*) in R (version 3.3.3); and associated scripts are available at (https://github.com/ctkremer/guppy).

From the set of candidate models fit to each separate fitness component and stream, we used AICc comparison to select the best model (Table S1, S4). For each of the four resulting models, we also: (i) obtained approximate 95% confidence bands, and (ii) estimated 95% confidence intervals for the value of the hybrid index where longevity or reproductive success was highest (i.e., the maxima of the quadratic function). Both of these analyses relied on simulating 10,000 new data sets based on the original model fits (using glmmTMB’s simulate function) and re-fitting the model to each data set. For each new fit, we calculated the position of the quadratic maxima; all values above 1 were rounded down (as hybrid index ranges from 0 to 1). Approximate 95% confidence intervals were then estimated as falling between the 2.5^th^ and 97.5^th^ quantiles of the resulting distribution. Confidence bands were obtained similarly, by constructing distributions of predicted regression lines at a range of hybrid indices.

### RADseq data collection

We collected genomic data using RADseq for a subset of 96 individuals collected before gene flow (20 from Caigual and 20 from Taylor collected in 2009) and again approximately 10 generations following the onset of gene flow (23 from Caigual and 23 from Taylor collected in 2012), as well as individuals from the downstream source population (10 from Guanapo collected in 2009). Genomic DNA for RADseq library preparation was purified from guppy scales using the DNeasy Blood & Tissue kit with an additional RNase A treatment following the manufacturer’s recommended protocols (Qiagen, CA, USA). Purified DNA was quantified using a Qubit and brought to equal concentration. We prepared RAD sequencing libraries for 96 individuals following the protocol of (*37*). RAD libraries were sequenced on an Illumina HiSeq sequencer at the University of Oregon with single-end 100 bp reads.

Raw sequence reads were demultiplexed using the process_radtags program in Stacks v.1.09 (*38*). Reads with sufficiently high sequencing quality that had the correct barcode and an unambiguous RAD site were retained. We used the guppy genome produced from a female guppy from the Guanapo source population as a reference sequence (version GCF_000633615.1_Guppy_female_1.0, (*22*)). Demultiplexed reads were aligned to the reference genome using GSnap (*32*). We required unique alignments, allowing for a maximum of five mismatches, the presence of up to two indels, and no terminal alignments. Aligned reads were analyzed using the ref_map.pl program in Stacks, which derived each locus from overlapping GSnap alignments to produce a consensus sequence. SNPs were determined and genotypes called using a maximum-likelihood statistical model implemented in Stacks. We removed four individuals that had greater than 50% missing loci. To include a locus in further analyses, we required it to be genotyped in at least 60% of individuals of each population. Only a single randomly chosen SNP per RAD locus was included. We removed loci with a minor allele frequency cutoff of less than 0.02 and loci below a log likelihood threshold of -30. At this point, we removed an additional six individuals that had an average read depth of less than ten. We did not test for linkage disequilibrium or conformation to Hardy-Weinberg proportions because we expected extensive physical linkage and non-random mating due to the extensive admixture between recipient and source populations.

### Detecting selection on locally adaptive variation

Briefly, we ran two ADMIXTURE (*33*) analyses: one for Caigual and one for Taylor. Each analysis estimated admixture proportions for the “pure” pre-gene flow population, the “pure” downstream source population, and the post-gene flow population. Using those outputs, we simulated many post-gene flow samples for each stream with the same admixture make-up and sampling noise. We also identified a set of loci that were strong candidates for alleles involved in (or linked to alleles involved in) local adaptation to the low predation (LP) headwater environment using two selection criteria: loci with (1) highly similar allele frequencies between Caigual and Taylor pre-gene flow populations, and (2) highly differentiated allele frequencies between both pre-gene flow populations and the downstream source (i.e., high *F*ST outliers). For each of these candidate loci, we then calculated an ancestry deviation, or the difference between the observed allele frequency in the post-gene flow population and the simulation-based expectation. This ancestry deviation measures the amount by which an allele’s frequency differs from the expectation in the direction of the inferred ancestral pre-gene flow frequency, suggesting directional selection for the locally adapted allele in the face of gene flow. To determine whether the observed deviations at our candidate SNP-set differed from the null, we matched each of our candidate alleles by frequency with a non-candidate locus, and compared the distribution of ancestry deviations in our candidate set to those of our frequency-matched non-candidate set. For example, if an observed candidate allele frequency in one of the post-gene flow populations was 0.8, it was matched with a non-candidate locus that had an expected post-gene flow allele frequency 0.8 (±0.05). The ancestry-polarized deviation for these frequency-matched loci is the difference between observed and expected shifts in allele frequencies, with a positive sign if the deviation was toward the ancestral LP frequencies inferred from the ADMIXTURE analyses, and negative otherwise. We then tested whether the distribution of deviations in candidate loci was different than the distribution of non-candidate loci. These methods are discussed in greater detail in Supplementary Appendix I. All candidate loci were used in a BLAST query against the Trinidadian guppy genome ((*34*); NCBI Poecilia reticulata Annotation Release 101; GCF_000633615.1).

## Acknowledgments

We thank P. Bois and the many field assistants who contributed to our guppy capture-mark-recapture study in Trinidad. We thank Cameron Ghalambor and David Reznick for their intellectual contribution to this work. Experimental methods were approved by the Colorado State University Institutional Animal Care and Use Committee (protocol no. 12-3818A). Collection permits were graciously provided by the Fisheries division of Trinidad’s Ministry of Food Production, Land and Marine Affairs. This is W.K. Kellogg Biological Station contribution no. xxxx.

## Author contributions

S.W.F., L.M.A., and W.C.F. designed the study; S.W.F., L.M.A., W.C.F., and P.E.S. performed the research; S.W.F., G.S.B., C.T.K., and P.E.S. analyzed data; all authors contributed to writing the paper.

## Data and materials availability

Data archiving to be completed prior to publication.

## Supplementary Materials

### Supplementary methods: Detecting selection on locally adaptive variation

In this section, we describe in detail the procedure used to identify the signature of selection for the maintenance of locally adaptive variation in the face of gene flow.

#### Logic

If pre-gene flow headwater populations were locally adapted to their headwater low predation (LP) habitat, and there has been selection for the maintenance of locally adapted variation in the face of gene flow, the signature of that selection would be greater-than-expected amounts of pre-gene flow ancestry in the post-gene flow headwater populations at and around the alleles involved in local adaptation. Identifying this signature can be difficult, as gene flow may homogenize even large allele frequency differences that have arisen between populations due to different directional selection. However, the sampling of individuals from the pre-gene flow headwater and source populations, as well as replication across two independent headwater populations, gives us power to determine whether there has been selection on locally adaptive variation.

#### Workflow

In this section, we briefly lay out the steps of our analysis, and in subsequent sections we go into greater depth on each step.

1. We ran ADMIXTURE to estimate admixture proportions for the post-gene flow headwater populations as well as allele frequencies in “pure” recipient pre-gene flow headwater and source populations.
2. Using those outputs, we simulated many post-gene flow stream samples with the same admixture make-up and sampling noise.
3. We then calculated an ancestry deviation for each locus in the dataset. This ancestry deviation measures the amount by which an allele’s frequency differs from the simulation-based expectation in the direction of the inferred ancestral headwater frequency, suggesting directional selection for the locally adapted allele in the face of gene flow.
4. Finally, we identified a set of loci that were strong candidates for alleles involved in local adaptation to the headwater environment prior to gene flow. To determine whether the observed deviations at our candidate SNP-set differed from the null, we matched each of our candidate alleles by frequency with a non-candidate locus, and compared the distribution of ancestry deviations in our candidate set to that of our frequency-matched non-candidate set.

We found that our candidate loci were significantly more “headwater” in their frequencies than their frequency-matched null set. This result supports the inference that headwater populations were locally adapted, and that there was selection for the maintenance of adaptive variation in the headwater habitats in the face of downstream gene flow.

#### Workflow (1): Clustering analyses

For each post-gene flow population, we generated a parametric null hypothesis for the frequency of each SNP, against which we could compare the observed frequencies to determine whether there was greater maintenance of local variation than we would expect. To generate this null for each stream, we first ran the model-based clustering method ADMIXTURE (*39*) on a dataset consisting of the two pre-gene flow populations (recipient and source) and the post-gene flow population, modeling individuals as draws from two discrete population clusters (*K*=2). We ran 10 replicate ADMIXTURE runs on each stream dataset, specifying a different random seed for each run to ensure that results were consistent across runs (they were).

We were interested in two quantities from each of these analyses: 1) the matrix of admixture proportions inferred for our post-gene flow populations (one admixture proportion per individual per cluster; Fig. S1); and 2) the vector of inferred allele frequencies in each canonical cluster (Fig. S2).

As expected, individuals from post-gene flow populations were of majority source ancestry in both streams (although to slightly differing degrees). The sampled source individuals were inferred to have some admixture with recipient populations in analyses in both streams, which is biologically plausible given the downstream direction of stream-flow.

In both streams, there were strong correlations between the estimated allele frequencies in the two inferred clusters and the sample allele frequencies observed in the pre-gene flow recipient and source populations. The allele frequencies estimated in “Cluster 2” in both runs, which corresponded most closely to the source (SGS) population, were quite similar to each other. The allele frequencies estimated for “Cluster 1” in both runs were very highly correlated with the sample frequencies observed in each respective pre-gene flow recipient population. The sample allele frequencies observed in the pre-gene flow recipient populations in each stream (NCA and NTY) were, for the most part, not strongly correlated with each other, presumably the result of the isolation they have experienced and the independent drift and/or selection they have undergone while isolated.

#### Workflow (2): Simulating admixed populations

We then used the admixture proportions and cluster allele frequencies estimated in ADMIXTURE to simulate admixed populations to match the observed post-gene flow populations. We had to use the ADMIXTURE-estimated frequencies, rather than observed pre-gene flow population frequencies, because some of our pre-gene flow samples were inferred to be admixed, and so did not offer a clear glimpse of the “pure” parental population frequencies. We describe the simulation procedure for a single stream below, using *w_ik_* to denote the estimated admixture proportion of the *i*th individual in the *k*th cluster, and *f_lk_*to denote the estimated allele frequency at the *l*th locus in the *k*th cluster.

To simulate a single haplotype in individual *i* within a stream, we randomly chose a fraction *w_i1_* of all genotyped loci to be of Cluster *1* ancestry, and assigned the remaining 1-*w_i1_* fraction of loci to be of Cluster 2 ancestry. At a given locus, we then simulated a haploid genotype as a Bernoulli draw with probability of success *f_lk_*; this step approximates the randomness of the sampling procedure used in the original genotyped dataset. We simulated two haplotypes across all loci for each individual, and repeated this procedure for each genotyped individual in the stream. We simulated 1000 replicate datasets for each stream. Note that we were only simulating data for the post-gene flow populations in our sample (Post-gene flow Caigual - PCA; Post-gene flow Taylor - PTY). Overall, we saw tight correlations between the simulated and observed data, with only 1.01% of loci falling outside the 95% quantile of simulations in Caigual, and 1.53% in Taylor.

#### Workflow (3): Calculating ancestry deviation

Using these simulations, we could calculate an estimated mean allele frequency at each locus, as well as an observed deviation from that expectation at each locus in each stream. The estimated mean allele frequency at a locus was defined as the mean of the simulated frequencies across all simulated replicates, and the deviation was the difference between the observed sample allele frequency in a post-gene flow population (either PCA or PTY) and the simulation mean. If our data were well described by the ADMIXTURE model, the distribution of deviations from the model-based expectation should have had mean zero and small variance. In practice, we saw that the distribution of deviations from expectation has mean 2.9 × 10^-3 and standard deviation 4.3 10^-2 in Caigual, and mean 2.6 × 10^-3 and standard deviation 4.2 × 10^-2 in Taylor.

We could further characterize the deviation at each locus by its direction: either toward the ancestral headwater frequency or the ancestral source frequency. To do this, we used the allele frequencies estimated in Clusters 1 and 2 from the ADMIXTURE analyses as the ancestral headwater and source population frequencies within each stream, respectively. We defined the ancestry-polarized deviation at a locus as positive when the difference between the observed sample frequency and the simulation-based expectation was in the direction of the ancestral pre-gene flow allele frequency, and negative when it was not (Fig. S3). If there was no difference between the observed and expected allele frequencies, the deviation was zero. We could then calculate the ancestry-polarized deviations across all loci in both streams (Fig. S3).

#### Workflow (4): Identifying excess pre-gene flow ancestry at putatively locally adapted loci

To detect a signal of excess pre-gene flow ancestry, we started by identifying alleles that matched our expectations for locally adapted loci. Alleles were included in our candidate list if the difference in sample frequencies between both pre-gene flow headwater populations was less than 0.1, and the frequency difference between each of the pre-gene flow headwater populations and the downstream source population was greater than 0.9. These are stringent criteria, so we undoubtedly have a high false negative detection rate, but as we are not interested in these loci individually, but rather in the signal of ancestry deviation aggregated across them, we feel the sacrifice in Type II error is worth the gains in Type I error.

In all, alleles at 146 loci met our criteria to be considered candidates, and we calculated the ancestry-polarized deviation for each of these loci. However, because the frequencies (both observed and expected) of these alleles affect the distribution of their deviations, it may not be appropriate to simply compare the distribution of ancestry-polarized deviations for these candidate loci to that of all loci, or all other loci.

Instead, we took the approach of comparing the distribution of ancestry-polarized deviations from the 146 candidate loci to that of a set of frequency-matched loci. To match by frequency at a locus, we chose another locus (that was not part of the candidate set) for which the mean simulated frequency fell within 0.05 of the observed post-gene flow allele frequency. We did this sampling without replacement, so that no two candidate loci were frequency-matched to the same locus.

We then calculated the distribution of ancestry-polarized deviation for this null set and compared it to that of our candidate loci to determine whether ancestry at the candidate loci was biased toward ancestral headwater frequencies. To assess significance, we used a one-tailed t-test, paired by locus, within each stream (Fig. 3C). We found that, in each stream, the frequencies of the candidate loci were significantly more “headwater” (pre-gene flow ancestry) than their frequency-matched null set. In Caigual, the mean deviation from prediction toward pre-gene flow upstream frequencies was 3.182e-2 for candidate loci and 6.566e-4 for non-candidate loci (one tailed paired t-test, *p* = 1.993e-7). In Taylor, the mean deviation from prediction toward pre-gene flow upstream frequencies was 2.002e-2 for candidate loci and 5.089e-3 for non-candidate loci (one tailed paired t-test, *p* = 3.724e-3).

## Supplementary Figures

**Fig. S1:**
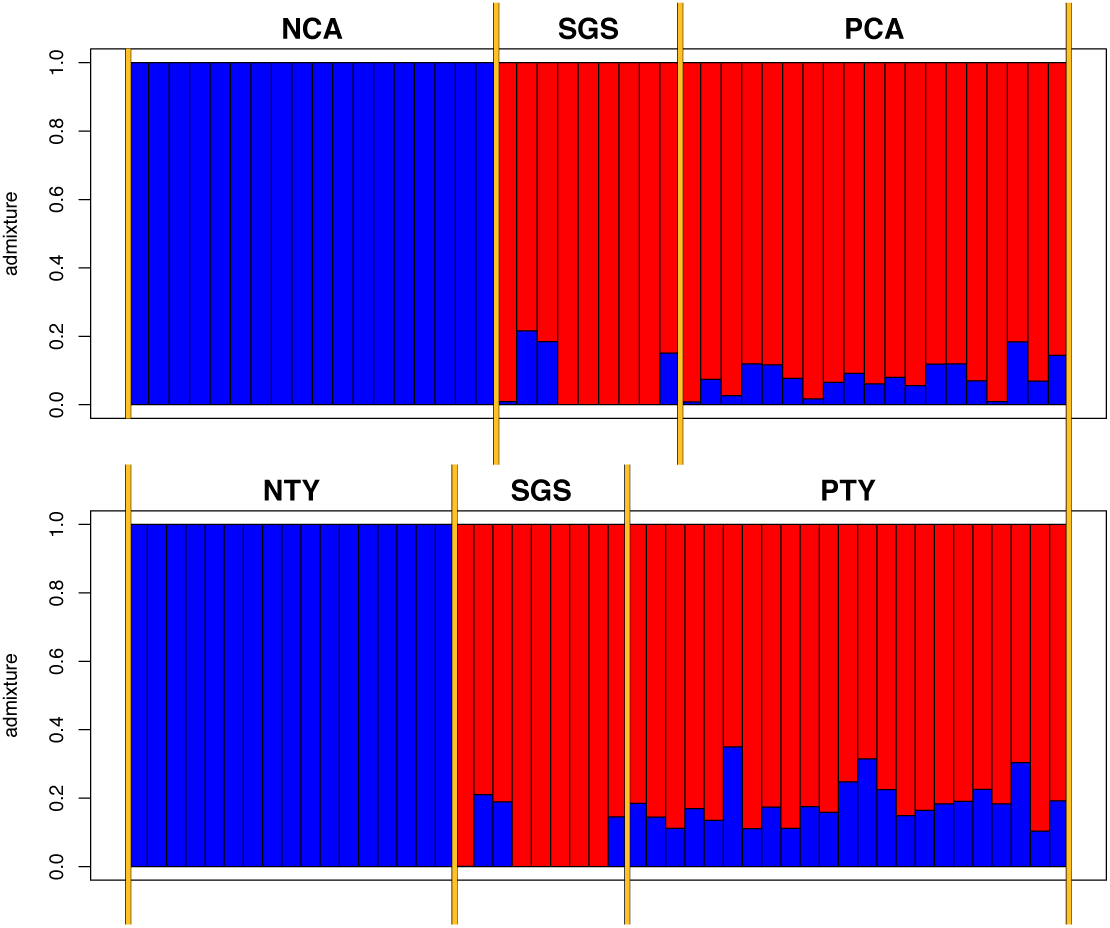
Structure plot of admixture proportions inferred by ADMIXTURE in each stream. Pre-gene flow recipient populations (pre-gene flow Caigual-NCA; pre-gene flow Taylor-NTY) were inferred to be of “pure” recipient ancestry, while the downstream source population (SGS) was inferred to be slightly admixed in analyses of both streams. Both post-gene flow populations were inferred to have majority SGS ancestry, although to slightly differing degrees.

**Fig. S2.**
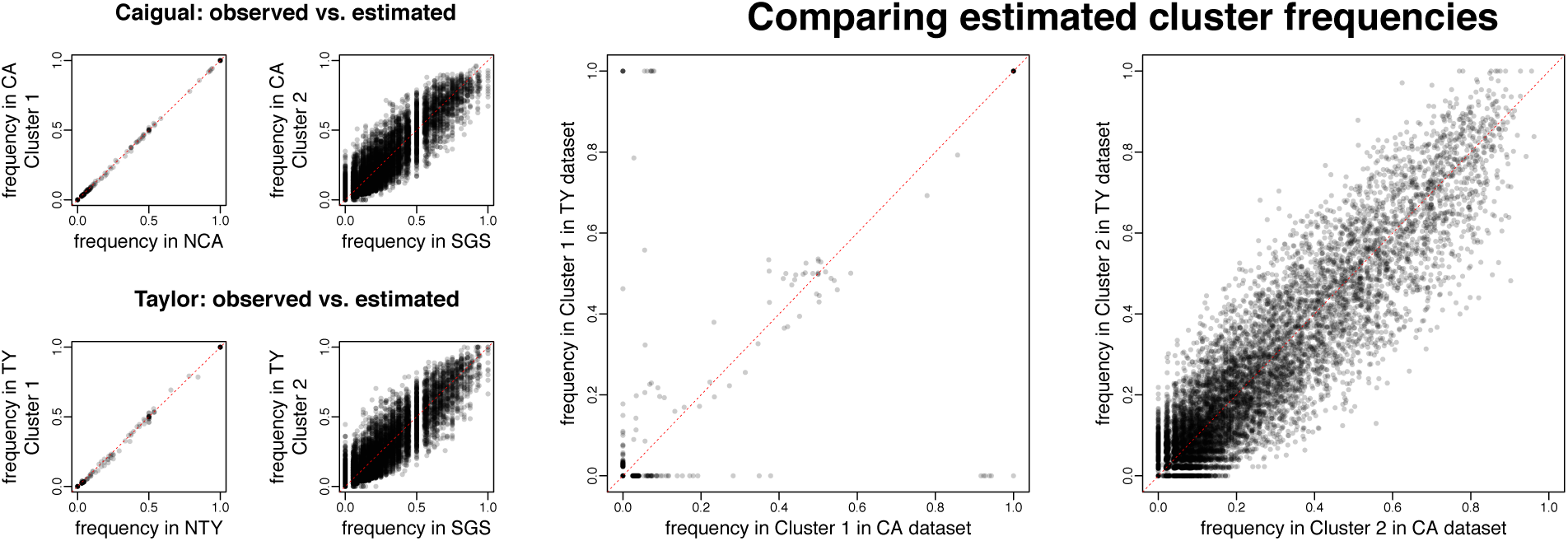
Observed and estimated allele frequencies for all SNPs. On the left, four small plots show the observed allele frequencies in the source population (SGS) and both pre-gene flow recipient populations (pre-gene flow Caigual - NCA; pre-gene flow Taylor - NTY) compared to the allele frequencies estimated by ADMIXTURE in each of the two clusters in each of the two analyses (one for each stream). On the right, two large plots compare the estimated allele frequencies in matched inferred clusters in the two runs (Caigual - CA; Taylor - TY).

**Fig. S3.**
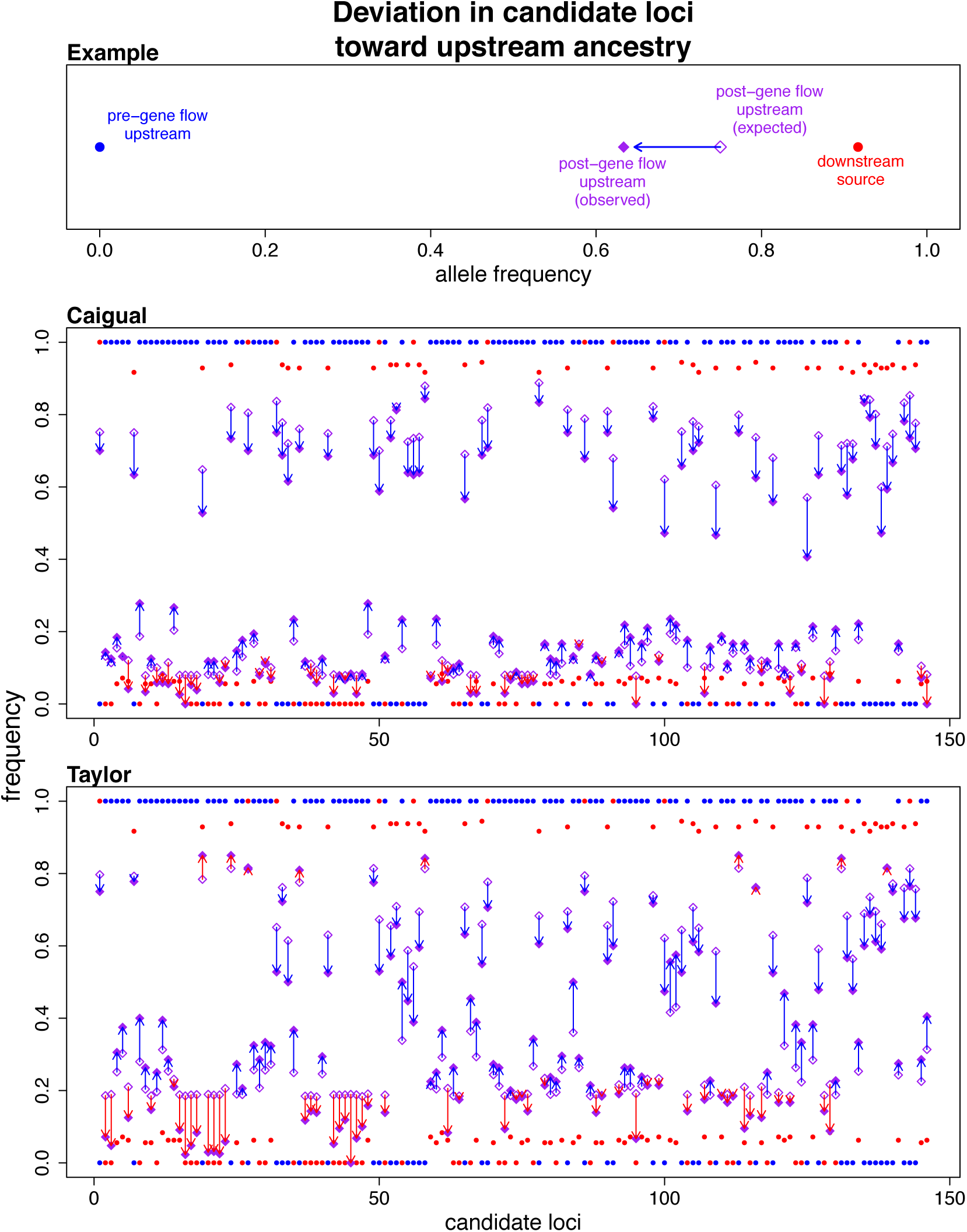
Deviation in candidate loci toward upstream ancestry. The top panel provides an example of ancestry-polarized deviation. In this example, the deviation between the observed and expected allele frequencies is in the direction of the ancestral pre-gene flow frequency, so the ancestry-polarized deviation would take a positive value. The middle panel is a plot of ancestry deviations across all candidate loci in the Caigual population. The bottom panel is a plot of ancestry deviations across all candidate loci in the Taylor population.

**Fig. S4.**
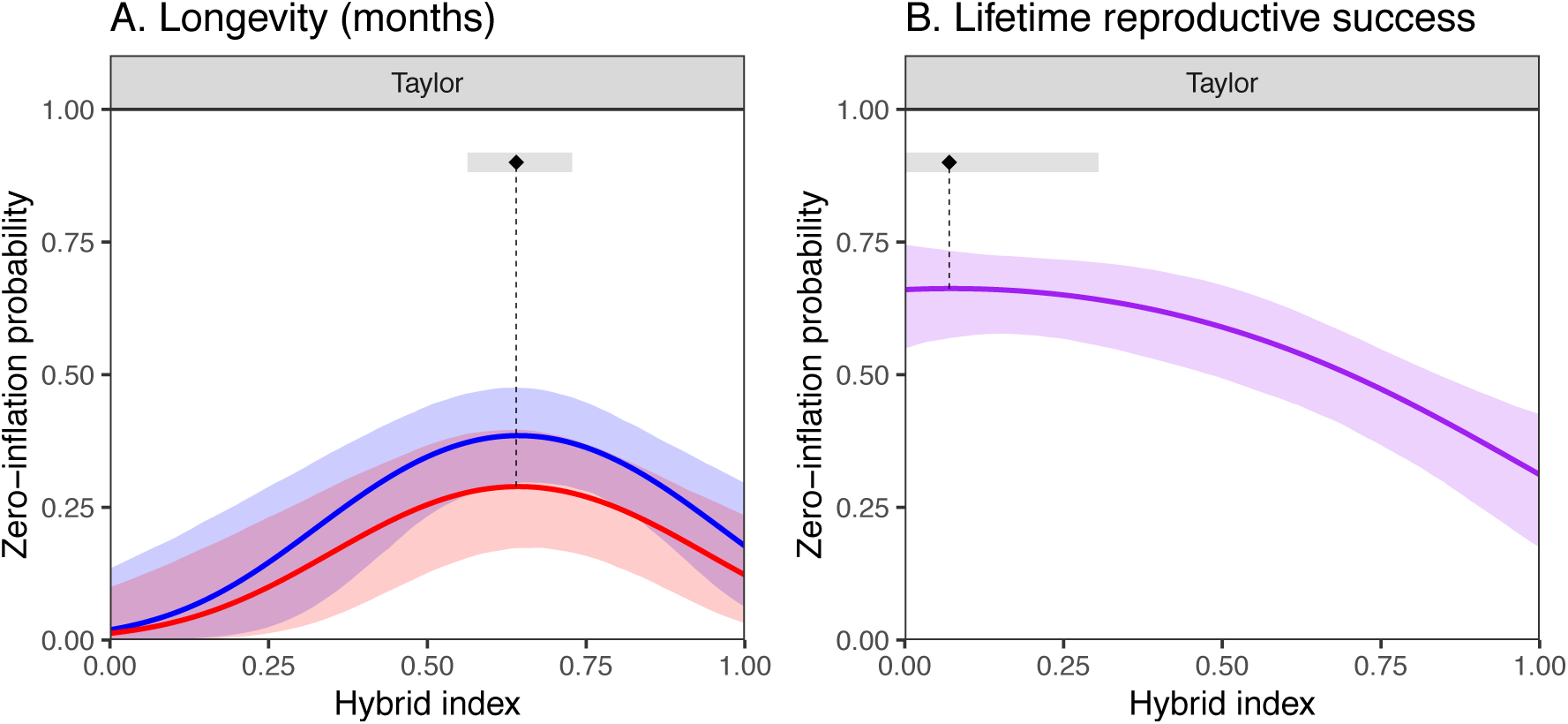
Zero inflation regressions. **A.** Fish with intermediate hybrid indices, especially females (blue), showed elevated zero inflation probabilities for longevity in Taylor. **B.** Zero inflation also occurred in the lifetime reproductive success of Taylor fish, independent of their sex. In both panels, bands around regression lines display approximate 95% confidence bands around regressions, obtained through simulation. Model details appear in Supplemental Tables S1-S5.

## Supplementary Tables

**Table S1.**
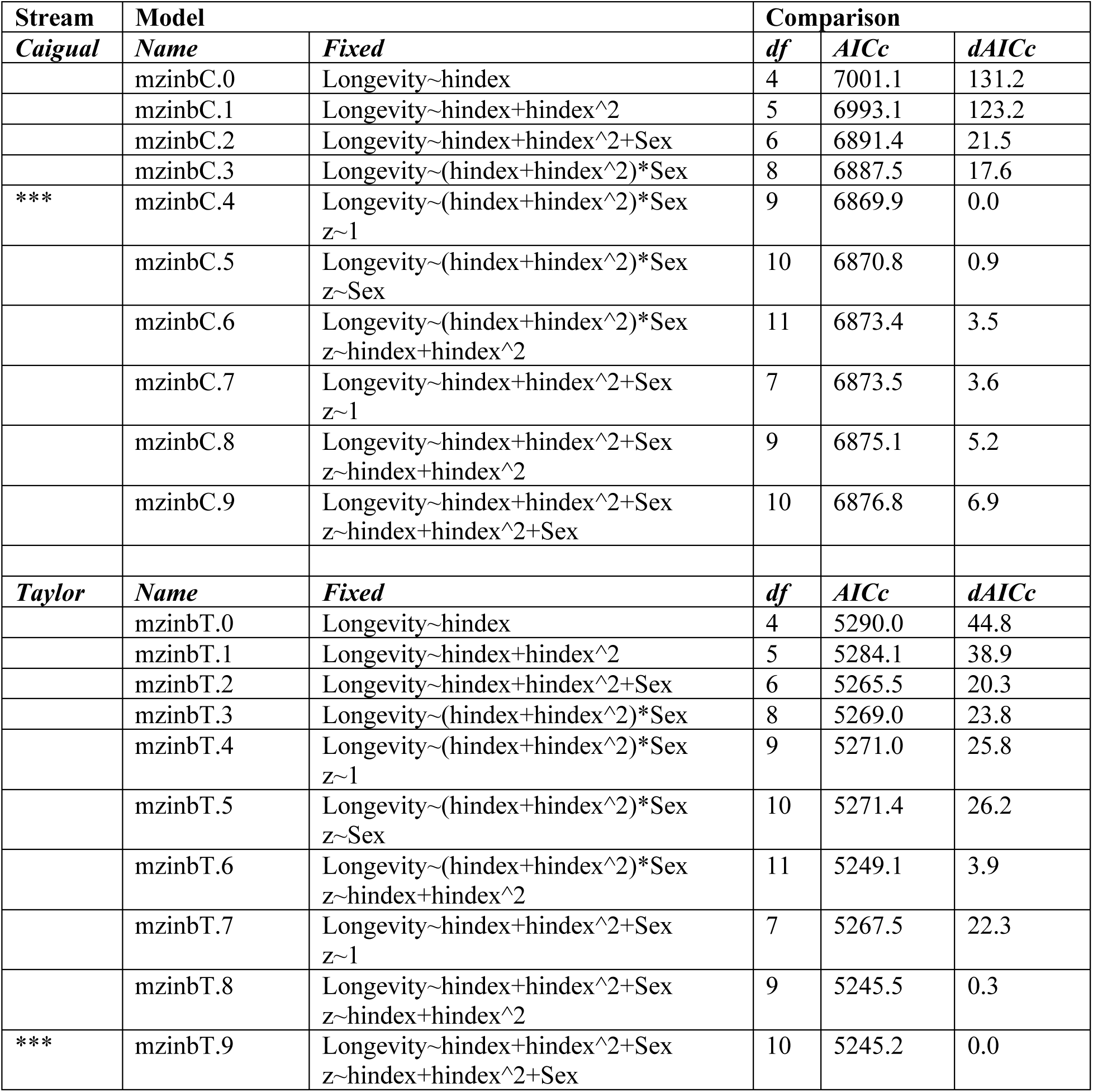
AICc comparison for Longevity models. All models included a random effect = 1|cohort. Other variables: z = zero-inflation sub-model; hindex = hybrid index; Sex = male/female. *** indicates the model selected as the best model for subsequent analyses.

**Table S2.** Detailed summaries of the best models of longevity by stream.

| Stream | Random effect | Variance | Std. Dev | Nobs | Nggroups |
| --- | --- | --- | --- | --- | --- |
| **Taylor**<br>(mzinbT.9 in Table S1) | Cohort | 0.3768 | 0.6139 | 1804 | 14 |
|  | **Negative binomial over-dispersion parameter** | 0.889 |  |  |  |
|  | **Negative binomial mean model** | **Estimate** | **Std. Error** | **z-value** | **Pr(>\|z\|)** |
|  | Intercept | -0.142 | 0.195 | -0.726 | 0.468 |
|  | hindex | 3.583 | 0.562 | 6.371 | <0.001 |
|  | hindex^2 | -2.382 | 0.537 | -4.435 | <0.001 |
|  | Sex(male) | -0.451 | 0.089 | -5.093 | <0.001 |
|  | **Zero-inflation model** | **Estimate** | **Std. Error** | **z-value** | **Pr(>\|z\|)** |
|  | Intercept | -3.940 | 1.062 | -3.708 | <0.001 |
|  | hindex | 10.795 | 3.101 | 3.481 | <0.001 |
|  | hindex^2 | -8.392 | 2.423 | -3.463 | <0.001 |
|  | Sex(male) | -0.432 | 0.303 | -1.425 | 0.154 |
| **Caigual**<br>(mzinbC.4 in Table S1) | **Random effect** | **Variance** | **Std. Dev** | **Nobs** | **Nggroups** |
|  | Cohort | 0.0450 | 0.212 | 1480 | 13 |
|  | **Negative binomial over-dispersion parameter** | 1.34 |  |  |  |
|  | **Negative binomial mean model** | **Estimate** | **Std. Error** | **z-value** | **Pr(>\|z\|)** |
|  | Intercept | 1.391 | 0.089 | 15.633 | <0.001 |
|  | hindex | 2.518 | 0.674 | 3.737 | <0.001 |
|  | hindex^2 | -2.134 | 1.018 | -2.096 | 0.036 |
|  | Sex(male) | -0.592 | 0.081 | -7.304 | <0.001 |
|  | hindex:Sex | -0.283 | 0.788 | -0.359 | 0.720 |
|  | hindex^2:Sex | -0.781 | 1.281 | -0.610 | 0.542 |
|  | **Zero-inflation model** | **Estimate** | **Std. Error** | **z-value** | **Pr(>\|z\|)** |
|  | Intercept | -1.907 | 0.203 | -9.383 | <0.001 |

**Table S3.** Position of quadratic maxima in longevity and lifetime reproductive success models (hybrid index values).

| **Fitness component** | **Stream** | **Term** | **Location** | **95% CI** |
| --- | --- | --- | --- | --- |
| Longevity | Caigual | Mean, Females | 0.590 | (0.428, 1.000) |
|  |  | Mean, Males | 0.383 | (0.302, 0.483) |
|  | Taylor | Mean, both | 0.752 | (0.667, 0.931) |
|  |  | Zero-inflation, both | 0.643 | (0.563, 0.728) |
| Lifetime reproductive success | Caigual | Mean | 0.824 | (0.663, 1.00) |
|  | Taylor | Mean | 0.678 | (0.585, 0.946) |
|  |  | Zero-inflation | 0.073 | (0.000, 0.309) |

**Table S4.**
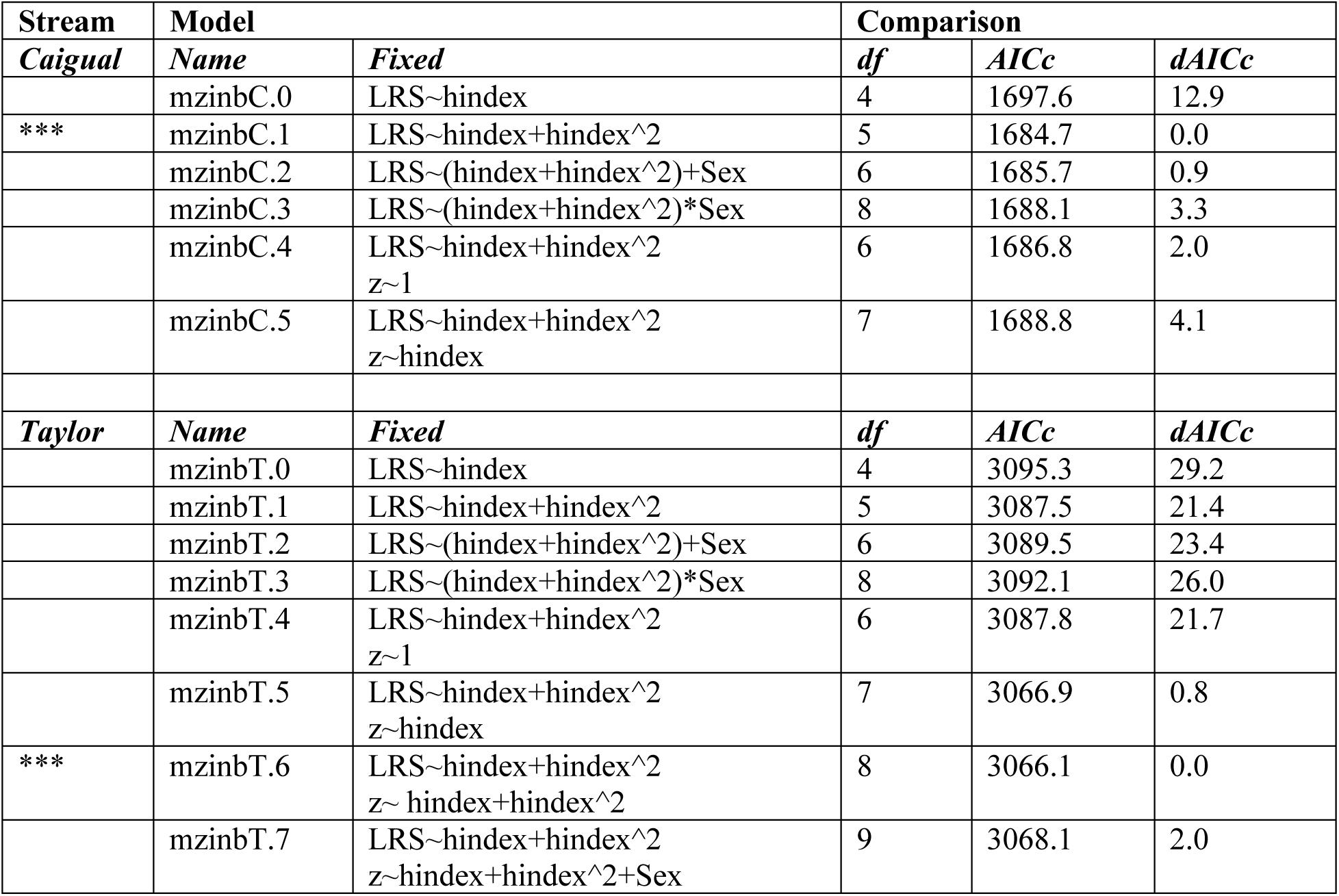
AICc comparison for lifetime reproductive success (LRS) models. All models included a random effect = 1|cohort. Other variables: z = zero-inflation sub-model; hindex = hybrid index; Sex = male/female. *** indicates the model selected as the best model for subsequent analyses.

**Table S5.** Detailed summaries of the best models of lifetime reproductive success by stream.

| **Stream** | **Random effect** | **Variance** | **Std. Dev** | **Nobs** | **Nggroups** |
| --- | --- | --- | --- | --- | --- |
|  | **Negative binomial over-dispersion parameter** | 0.65 |  |  |  |
|  | **Negative binomial conditional mean model** | **Estimate** | **Std. Error** | **z-value** | **Pr(>\|z\|)** |
|  | Intercept | 0.586 | 0.257 | 2.281 | 0.023 |
|  | hindex | 3.706 | 0.889 | 4.169 | <0.001 |
|  | hindex^2 | -2.732 | 0.828 | 3.300 | <0.001 |
|  | **Zero-inflation model** | **Estimate** | **Std. Error** | **z-value** | **Pr(>\|z\|)** |
|  | Intercept | 0.664 | 0.243 | 2.728 | 0.006 |
|  | hindex | 0.250 | 0.983 | 0.255 | 0.799 |
|  | hindex^2 | -1.707 | 1.037 | -1.646 | 0.100 |
| ***Taylor***<br>(mzinbT.6 in Table S5) | Cohort | 0.262 | 0.512 | 1110 | 11 |
| ***Caigual***<br>(mzinbC.1 in Table S5) | **Random effect** | **Variance** | **Std. Dev** | **Nobs** | **Nggroups** |
|  | Cohort | 1.742 | 1.32 | 702 | 10 |
|  | **Negative binomial over-dispersion parameter** | 0.453 |  |  |  |
|  | **Negative binomial mean model** | **Estimate** | **Std. Error** | **z-value** | **Pr(>\|z\|)** |
|  | Intercept | -1.209 | 0.438 | -2.757 | 0.006 |
|  | hindex | 9.691 | 1.100 | 8.807 | <0.001 |
|  | hindex^2 | -5.878 | 1.411 | -4.166 | <0.001 |

